# Distinct SARS-CoV-2 Antibody Reactivity Patterns in Coronavirus Convalescent Plasma Revealed by a Coronavirus Antigen Microarray

**DOI:** 10.1101/2020.09.16.300871

**Authors:** Rafael Assis, Aarti Jain, Rie Nakajima, Algis Jasinskas, Saahir Khan, Larry J. Dumont, Kathleen Kelly, Graham Simmons, Mars Stone, Clara Di Germanio, Michael Busch, Philip L. Felgner

## Abstract

A coronavirus antigen microarray (COVAM) was constructed containing 11 SARS-CoV-2, 5 SARS-1, 5 MERS, and 12 seasonal coronavirus recombinant proteins. The array is designed to measure immunoglobulin isotype and subtype levels in serum or plasma samples against each of the individual antigens printed on the array. We probed the COVAM with COVID-19 convalescent plasma (CCP) collected from 99 donors who recovered from a PCR+ confirmed SARS-CoV-2 infection. The results were analyzed using two computational approaches, a generalized linear model (glm) and Random Forest (RF) prediction model, to classify individual specimens as either Reactive or Non-Reactive against the SARS-CoV-2 antigens. A training set of 88 pre-COVID-19 specimens (PreCoV) collected in August 2019 and102 positive specimens from SARS-CoV-2 PCR+ confirmed COVID-19 cases was used for these analyses. Results compared with an FDA emergency use authorized (EUA) SARS-CoV2 S1-based total Ig chemiluminescence immunoassay (Ortho Clinical Diagnostics VITROS® Anti-SARS-CoV-2 Total, CoV2T) and with a SARS-CoV-2 S1-S2 spike-based pseudovirus micro neutralization assay (SARS-CoV-2 reporter viral particle neutralization titration (RVPNT) showed high concordance between the 3 assays. Three CCP specimens that were negative by the VITROS CoV2T immunoassay were also negative by both COVAM and the RVPNT assay. Concordance between VITROS CoV2T and COVAM was 96%, VITROS CoV2T and RVPNT 93%, and RVPNT and COVAM 95%. The discordances were all weakly reactive samples near the cutoff threshold of the VITROS CoV2T immunoassay. The multiplex COVAM allows CCP to be grouped according to antibody reactivity patterns against 11 SARS-CoV-2 antigens. Unsupervised K-means analysis, via the gap statistics, as well as hierarchical clustering analysis revealed 3 main clusters with distinct reactivity intensities and patterns. These patterns were not recapitulated by adjusting the VITROS CoV2T or RVPNT assay thresholds. Plasma classified according to these reactivity patterns may be better associated with CCP treatment efficacy than antibody levels alone. The use of a SARS-CoV-2 antigen array may be useful to qualify CCP for administration as a treatment for acute COVID-19 and to interrogate vaccine immunogenicity and performance in preclinical and clinical studies to understand and recapitulate antibody responses associated with protection from infection and disease.

## Background

Following exposure and recovery from SARS-CoV-2 infection, convalescent patients develop antigen specific adaptive T-and B-cell immune responses including binding and neutralizing antibodies (Ab). These immune responses may prevent reinfection or blunt the clinical consequences of future infectious exposures to the virus. Administration of COVID-19 convalescent plasma (CCP) from recovered patients is being employed for therapeutic use based on the belief that factors, including neutralizing Ab against SARS-CoV-2, present in the plasma may inhibit virus replication and improve patient outcomes.

Numerous clinical trials are underway aimed at understanding how best to administer CCP and to test the hypothesis that it is an effective treatment. A significant clinical trial involving more than 35,000 CCP treated COVID-19 patients has shown an efficacy signal from plasma with elevated Ab levels in patients treated within a few days after symptom onset [1]. Previous efforts to use convalescent plasma for treatment of infectious diseases have produced mixed results, possibly associated with differences in the antibody profile from different donors. Differences in the breadth and level of SARS-CoV-2-specific Ab in CCP may correlate with therapeutic efficacy. Consequently, investigated the performance of a Coronavirus Antigen Microarray (COVAM) in order to characterize and classify Ab reactivity in CCP before it is administered to patients.

COVAM is a multiplex assay platform for high-throughput serological studies. The microarrays are produced by printing validated and purified recombinant antigens on nitrocellulose-coated slides. The COVAM has 11 SARS-CoV-2 antigens, 5 SARS-1, 5 MERS and 12 seasonal coronavirus antigens, as well as 35 antigens from 5 other viruses that cause acute respiratory infections [10,11]. A complete list of the COVAM antigens can be found in Supplementary Table 1. The arrays are designed to determine the Ab profile in serum or plasma samples to confirm prior exposure following suspected infection with the viruses and to monitor Ab changes over time. The arrays are designed for high throughput and low-cost testing and hence are suitable for epidemiology studies as a serologic surveillance tool to determine the prevalence and levels of Abs indicating viral exposure within individuals over the course of an epidemic.

Here we probed the COVAM with CCP collected from 99 donors between 4/18/2020 – 5/6/2020 from 8 regions across the US. Although the donors were all recovered SARS-CoV-2 PCR+ COVID-19 cases, their Ab response profiles against 11 SARS-CoV-2 fell into distinct groups with different Ab levels and breadth of the responses particularly to 4 SARS-CoV-2 antigens. If these classification groups correlate with CCP efficacy, a SARS-CoV-2 antigen microarray may be useful to qualify CCP for treatment of acute COVID-19.

## Methods

### Specimen Testing on Coronavirus Antigen Microarray

The COVAM slides were probed with human plasma and analyzed as described elsewhere (Jain *et. al*.,2016; Nakajima *et. al*., 2018 and Khan *et al*., 2019). Briefly, the specimens were diluted 1:100 in 1X Protein Array Blocking Buffer (GVS Life Sciences, Sanford, ME), transferred to the slides and incubated overnight at 4°C. The slides were then washed 3 times for 5 minutes each with t-TBS buffer (20 mM Tris-HCl, 150 mM NaCl, 0.05% Tween-20 in ddH_2_O adjusted to pH 7.5 and filtered) at room temperature (RT). Then, anti-human IgG and anti-human IgA secondary Ab were added to each pad at a dilution of 1:100 (in Protein Array blocking Buffer) and incubated for 2h at RT under agitation. Pads were then washed with t-TBS 3 times for 5 minutes each and dried. The slides were imaged using ArrayCam imager (Grace Bio-Labs, Bend, OR). In order to measure non-specific binding of the secondary Ab, pads were incubated without the previous addition of human sera. The data acquisition and spot quantification were performed using the Scan ArrayExpress (V 3.0, PerkinElmer) software [2, 8-9].

### Specimen Sources and prior Characterization

In order to build the prediction models, a collection of samples with known exposure status to SARS-CoV-2 was used as a training set. This set was composed of 88 PreCoV specimens derived from frozen plasma components collected by Vitalant in July-September 2019, and 102 specimens from PCR confirmed cases was used for these analyses. These confirmed positive cases include 45 serum samples from the University of California Irvine collected between 7 and 25 days after the symptom onset; 30 samples from the University of California San Francisco collected between 2 and 38 days after the symptoms onset; 13 samples from Basel collected between 13 and 50 days after the symptoms onset and 14 samples from Ortho Clinical Diagnostics collected between 7 and 22 days after the symptoms onset. The CCP samples were a collection of 99 SARS-CoV-2 convalescent plasma samples from the Vitalant system collected between 4/18/2020 – 5/6/2020 from 8 regions across the US.

Neutralization titers were measured as 50% neutralization (NT-50) by endpoint titration using a recombinant viral-particle neutralization test (RVPNT) and a cutoff equal to 1:40. The *VITROS CoV2T chemiluminescent immunoassay assay* was performed using either serum or plasma samples from Vitalant Research Institute (San Francisco, CA) according to the manufacturer instructions [3]. The test targets the spike protein and applies a predefined threshold value of 1.0 signal-to-cutoff (S/C) for IgG seropositivity but has a broad dynamic range with S/C values as high as 1000.

## Data Analysis

All data analysis was performed in the R programing environment (Version 4.0.2).

### Quantile normalization

After data acquisition, data normalization was performed by the Quantile Normalization method using the “normalize.quantiles.use.target” function of the “preprocessCore” package (Version 1.50.0). As reference for normalization, a collection of known positive and known negative samples (training set comprised of serial samples from recovered COVID-19 patients and plasma collected in the July-September 2019, respectively) was used.

### Cluster analysis

To investigate the different reactivity profiles, the data were clustered and divided based on the reactivity to the 11 SARS-CoV-2 antigens. For the clustering, first the optimal number of clusters was estimated by the gap statistics on the K-means clustering analysis (“fviz_nbclust” function from the factoextra package (version 1.0.7). Then, a Hierarchical Clustering analysis (“hclust” function, method “ward.D2”, from the “stats” package, version 4.0.2) was performed and the dendrogram cut in order to obtain a number of cluster defined by the previous calculated gap statistics. The assigned groups were obtained by cutting the dendrogram to obtain K = n groups, where n is the optimal number of clusters obtained from the gap statistics analysis.

Principal component Analysis (PCA) was also performed (“prcomp” function from the “stats” package version 4.0.0 as well as the “fviz_nbclust” function from the “factoextra” package, version 1.0.7)

### Reactivity classification

With goal of predicting exposure to SARS-CoV-2, the samples were first classified based on their overall reactivity profile. The prediction was performed using two main computational methods, a logistic regression model and a random forest prediction model.

For the logistic regression, a generalized linear model (“logit” family) was first generated (“glm” function from the “stats” package version 4.0.2) using the training set of. Then a receiver operating characteristic curve (ROC) was generated in order to obtain all the curve coordinates. This allows estimation of the specificity and sensitivity for each point of the curve and therefore defines an optimal cutoff point of the regression analysis fitted values. For the predictions described in this work, the model was built using four antigens: SARS-CoV-2 S1, SARS-CoV-2 S1.HisTag, SARS-CoV-2.S1.RBD, SARS.CoV.2.S1+S2. The test samples are then submitted to the logistic regression analysis and the fitted values compared to the defined cutoff for classification (positive or negative).

For the random forest analysis, like the logistic regression method, a model fit is generated (random forest version 4.6-14). For the random forest model, seven antigens were used: SARS-CoV-2 NP, SARS-CoV-2 S1.HisTag, SARS-CoV-2 Spike.RBD.rFc, SARS.CoV.2.S1.mFcTag, SARS.CoV.2.S1.RBD, SARS.CoV.2.S1+S2, SARS.CoV.2.S2.

## Results

### COVAM, RVPNT and VITROS CoV2T chemiluminescent immunoassay assay concordance

The COVAM has 11 SARS-CoV-2 antigens, 5 SARS-1, 5 MERS and 12 seasonal coronavirus antigens. The complete list of the COVAM antigens, in the order displayed in the figures, is available in **Supplementary Table 1**. We probed COVAM with CCP collected from 99 US PCR confirmed COVID-19 plasma donors who recovered from the infection one to two months prior to CCP collection. The multiplex COVAM results were analyzed using two computational approaches, either a generalized linear model (glm) or Random Forest (RF) to classify individual specimens as either Reactive or Non-Reactive against the SARS-CoV-2 antigens. The binary prediction results were compared with the FDA EUA VITROS CoV2T chemiluminescence immunoassay from Ortho (reactive: S/C≥1.0) and with an RVPNT assay (reactive: neutralization titer with >5) inhibition of infection [NT50] > 40) developed at Vitalant and tabulated in **Table 1**.

| CCP # | VITROS CoV2T S/C Ratio | VITROS CoV2T Interpretation | RVPNT NT50 | RVPNT Interpretation | COVAM RF probability | COVAM RF Interpretation | Concordance |  |  |
| --- | --- | --- | --- | --- | --- | --- | --- | --- | --- |
|  |  |  |  |  |  |  | CoV2T vs RVPNT | CoV2T vs COVAM | RVPNT vs COVAM rf |
| 1 | 0.01 | Non-Reactive | <40 | Non-Reactive | 0.51 | Non-Reactive | 1 | 1 | 1 |
| 2 | 0.02 | Non-Reactive | <40 | Non-Reactive | 0.22 | Non-Reactive | 1 | 1 | 1 |
| 3 | 0.31 | Non-Reactive | <40 | Non-Reactive | 0.33 | Non-Reactive | 1 | 1 | 1 |
| 4 | 1.39 | Reactive | 280 | Reactive | 0.99 | Reactive | 1 | 1 | 1 |
| 5 | 1.45 | Reactive | 146 | Reactive | 0.47 | Non-Reactive | 1 | 0 | 0 |
| 6 | 2.14 | Reactive | 302 | Reactive | 1.00 | Reactive | 1 | 1 | 1 |
| 7 | 2.77 | Reactive | 4171 | Reactive | 1.00 | Reactive | 1 | 1 | 1 |
| 8 | 4.24 | Reactive | <40 | Non-Reactive | 0.72 | Reactive | 0 | 1 | 0 |
| 9 | 4.48 | Reactive | <40 | Non-Reactive | 0.63 | Reactive | 0 | 1 | 0 |
| 10 | 5.21 | Reactive | 617 | Reactive | 1.00 | Reactive | 1 | 1 | 1 |
| 11 | 5.36 | Reactive | 132 | Reactive | 0.73 | Reactive | 1 | 1 | 1 |
| 12 | 5.62 | Reactive | 3583 | Reactive | 1.00 | Reactive | 1 | 1 | 1 |
| 13 | 6.22 | Reactive | 98 | Reactive | 0.79 | Reactive | 1 | 1 | 1 |
| 14 | 7.54 | Reactive | <40 | Non-Reactive | 0.38 | Non-Reactive | 0 | 0 | 1 |
| 15 | 9.37 | Reactive | <40 | Non-Reactive | 0.24 | Non-Reactive | 0 | 0 | 1 |
| 16 | 9.45 | Reactive | <40 | Non-Reactive | 0.78 | Reactive | 0 | 1 | 0 |
| 17 | 9.80 | Reactive | 133 | Reactive | 1.00 | Reactive | 1 | 1 | 1 |
| 18 | 10.30 | Reactive | <40 | Non-Reactive | 0.26 | Non-Reactive | 0 | 0 | 1 |
| 19 | 11.00 | Reactive | 6910 | Reactive | 1.00 | Reactive | 1 | 1 | 1 |
| 20 | 12.40 | Reactive | <40 | Non-Reactive | 0.66 | Reactive | 0 | 1 | 0 |
| 21-99 | 12.9 - 436 | Reactive | 132 | Reactive | 0.69-1.0 | Reactive | 79 | 79 | 79 |
| % concordance |  |  |  |  |  |  | 93% | 96% | 95% |

Compared to the VITROS CoV2T assay, there is 93%, and 96% concordance with the RVPNT, and COVAM predictions, respectively. Three specimens that were negative by the VITROS CoV2T assay were also negative by COVAM and RVPNT assays. Ninety-one specimens (92%) are completely concordant among all comparisons, including 88 seropositive by all tests and 3 seronegative by all tests. All of the discordances are found in 8 samples that showed low positive titer on the VITROS CoV2T immunoassay. Among these 8 samples, 7 were considered non-reactive on the RVPNT test and 4 considered non-reactive on the COVAM prediction (using a 60% RF probability cutoff).

The COVAM data are summarized in the heatmap in **Figure 1**. There are 99 specimen IDs along the x-axis and 65 antigens along the y-axis (Supplementary Table 1). The antigens are grouped according to the virus from which they are expressed. The top 11 antigens are from SARS-CoV-2 and the quantitative COVAM IgG Ab results from these antigens were used to cluster the specimens into 3 groups as shown in the dendrogram on top of the heatmap (**Figure 1**).

**Figure 1.**
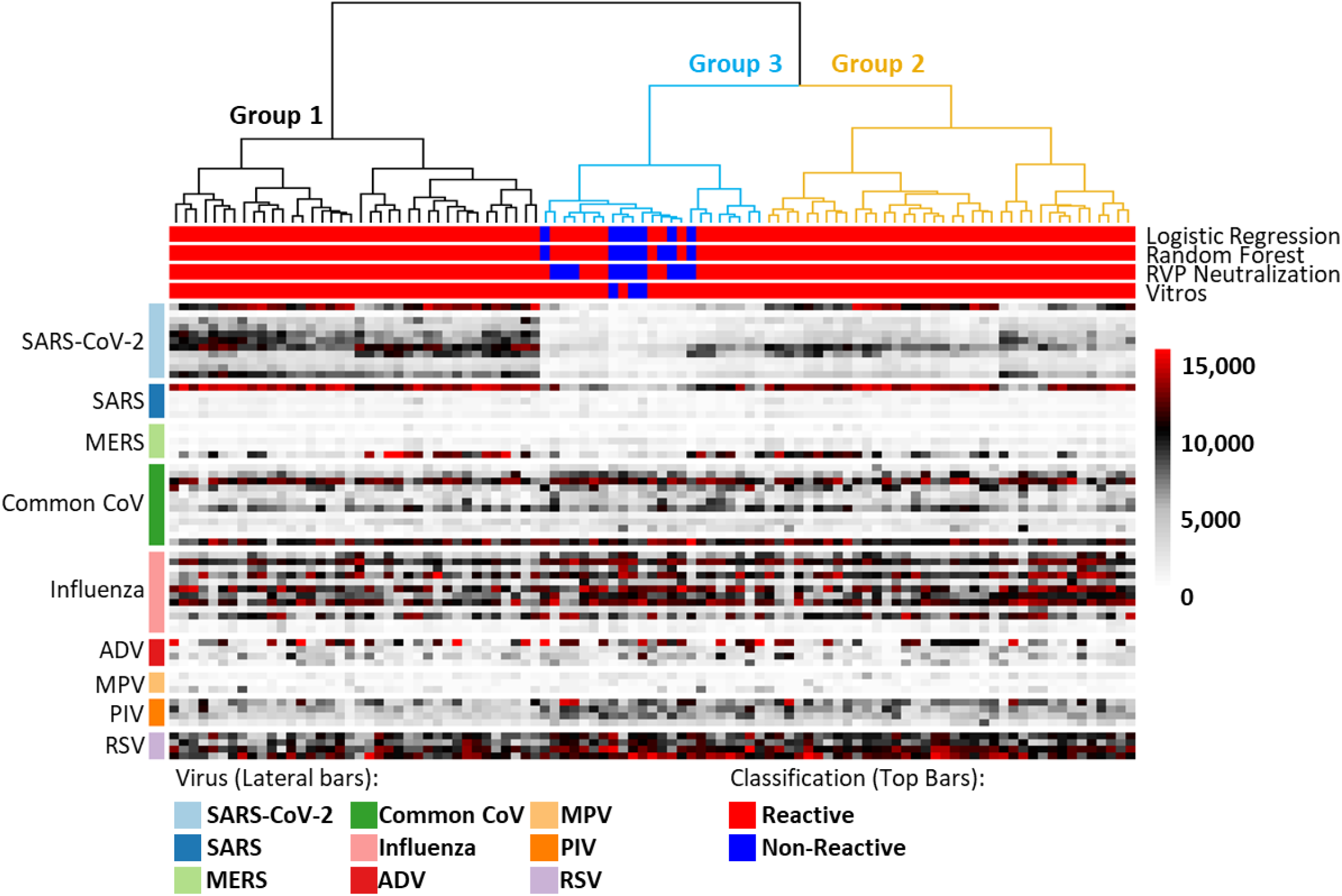
IgG COVAM reactivity heatmaps of 99 sera from recovered coronavirus convalescent cases. The heatmaps show the IgG reactivity levels of patients to antigens printed onto the COVAM array. Each column is the representation of a specific specimen while each row is represented as the mean reactivity of 4 replicates of each antigen. The COVAM array shows clear IgG reactivity to SARS-CoV-2 antigens that are clustered into 3 groups. A complete list of the antigens, in the order displayed, is available in the supplemental table 1.

The vsn normalized reactivities of individual antigens printed on the COVAM array can be correlated with the VITROS CoV2T immunoassay S/C values as well as the RVPNT titers. The results of correlation analysis for the S1 Spike antigen in the COVAM array, are shown in **Figure 2**. The normalized signal intensity on the COVAM for each antigen is plotted in a scatterplot against the either the VITROS CoV2T S/C levels or the RVPNT NT50 titers. The COVAM reactivity of this antigen correlates to the VITROS CoV2T S/Cs and to the RVPNT NT50 titers with Spearman’s r = 0.65 and r = 0.655, respectively. A summary of Spearman r values for each of the 11 SARS-CoV-2 COVAM are plotted in **Figure 2b**. Antigens that contain the RBD domain and lack the S2 domain correlate best. The COVAM reactivity against the nucleocapsid protein NP and the S2 domain of spike do not correlate with the VITROS S1 based CoV2T or the RVPNT assays, which is reasonable since Ab-mediated neutralization of virus entry is primarily based on blocking interactions between the RBD domain of S1 and ACE-2 receptors expressed on target cells in the assay.

**Figure 2.**
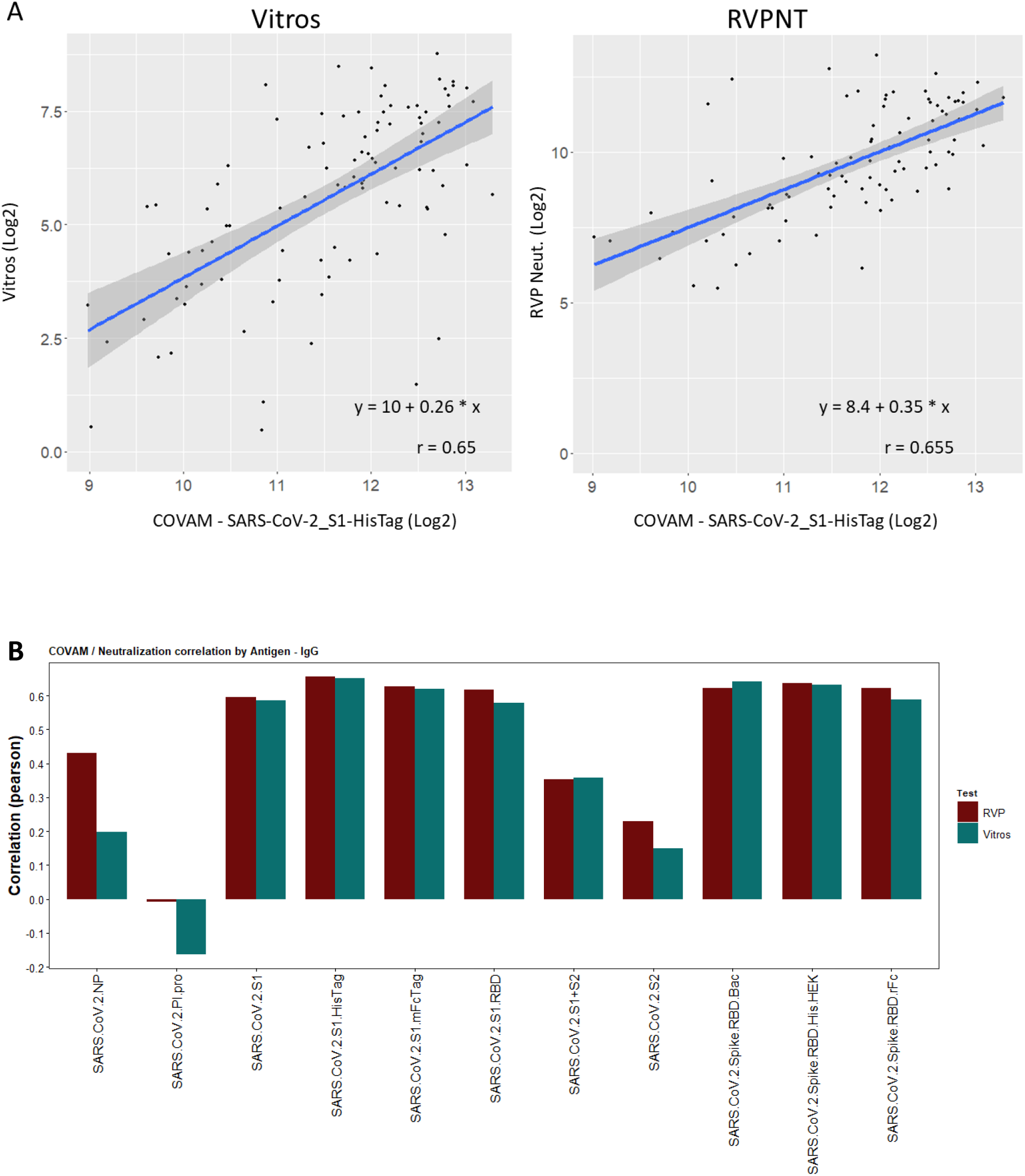
Correlation of the RVPNT assay NT50 values or VITROS CoV2T assay S/C values with the COVAM reactivity (RF values) to SARS-CoV-2 antigens. Panel A, scatterplots representative of the correlation between the vsn normalized COVAM reactivities and the RVPNT or VITROS CoV2T assays to the S1 (HisTag) antigen. Panel B, the Spearman correlation (r) of the vsn normalized COVAM reactivities and RVPNT or VITROS CoV2T Assay to the COVAM SARS-CoV-2 antigens

### CCP specimens cluster into 3 distinct groups according to Ab levels against SARS-CoV-2 antigens

Although there is concordance in the binary predictions of the 3 assays, the multiplex COVAM assay allows for a multivariant cluster analysis metric to classify CCP into different groups based on their reactivity patterns against several antigens. The VITROS CoV2T or RVPNT assays produce a single value for each specimen and a binary (reactive or non-reactive) result when that value is above or below a reactivity threshold, with quantitation of Ab intensity above the S/C = 1.0 and NT50 = 40 categorical thresholds. The COVAM interrogates IgG levels against 11 SARS-CoV-2 antigens and uses a computational algorithm to determine a binary outcome by comparing the results to the training set of COVID positive and negative controls.

Unsupervised cluster analysis can sort the specimens according COVAM antigen specific IgG reactivity patterns into at least 3 distinct groups based on the level of antibodies against a collection of NP, Spike, S1, S2 and RBD antigens (**Figure 1**). Group 1 specimens have high Ab levels against many of the SARS-CoV-2 antigens, most markedly high reactivity to nucleocapsid protein (NP), S1 domain of spike protein, and the full spike S1+S2 protein. Group 3 specimens have very low antibody levels to NP and S1 compared to the Group 1 specimens. The Group 2 specimens show an intermediate reactivity level against spike antigens. The horizontal bars on top of the heatmap indicate reactive (red) and non-reactive (blue) predictions for each specimen for the VITROS CoV2T, RVPNT and COVAM assays. All of the predicted non-reactive specimens from all three assays are in group 3. All of the group 1 and 2 specimens are reactive and concordant by all 3 assays.

The results in **Figure 3** show the IgG heatmap clustered (k-means), using the reactivity of the SARS-CoV-2 antigens, into these 3 groups. PCA analysis shows 3 distinct groups and the bar graphs confirm that Ab levels against each antigen differ significantly between each group. The heatmap and its dendrogram (**Figure 3**) indicate that the specimens in each group can be further split into subgroups. **Figure 4** separates Group 1 specimens into 2 subgroups (1.1 and 1.2) evident from the heatmap (**Figure 4a**) and confirmed by the PCA (**Figure 4c**). The bar graph (**Figure 4b**) highlights a significant difference between subgroups 1.1 and 1.2 in reactivity against antigen S2 which is elevated in Group 1.2. **Figure 5** separates Group 2 specimens into 2 subgroups (2.1 and 2.2) evident from the heatmap (**Figure 5a**) and confirmed by PCA (**Figure 5c**). The bar graph (**Figure 5b**) highlights a significant difference between subgroups 2.1 and 2.2 against the nucleocapsid protein and S2 which are both elevated in Group 2.2). **Figure 6** separates Group 3 specimens into 2 groups (3.1 and 3.2) evident from the heatmap (**Figure 6a**) and confirmed by PCA (**Figure 6c**). From the bar graph and heat map it is evident that the reactivity of group 3 specimens is less than that of the other two groups, but the subgroup 3.2 is particularly lacking in antibodies to SARS-CoV-2 antigens.

**Figure 3.**
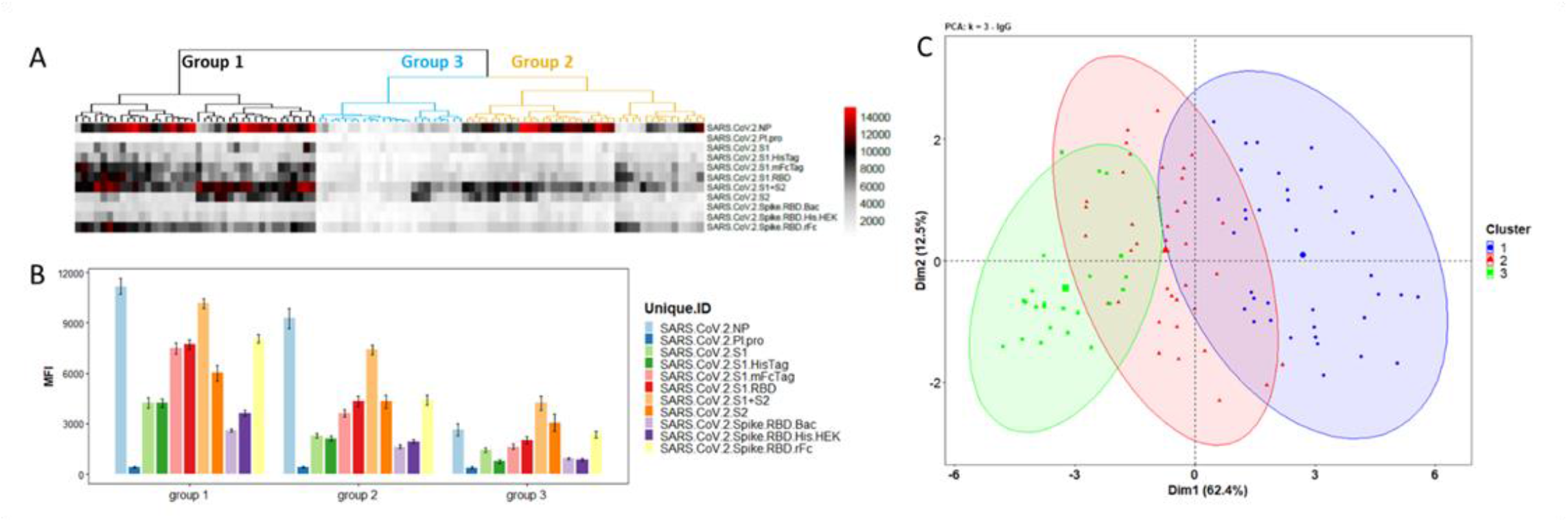
Clustering analysis of the IgG reactivity to the COVAM SARS-CoV-2 antigens. On A, heatmap showing the reactivity to the SARS-CoV-2 antigens. Samples were clustered using the hierarchical Clustering analysis. The dendrogram was cut (and color coded) to a final cluster number equal 3. On B, a bar graph showing the mean reactivity and the standard error of each cluster to each individual SARS-CoV-2 antigen. On C, principle component analysis (PCA) showing the spatial distribution of the samples for the first and second principal components that explain, combined, over 74% of the variance.

**Figure 4.**
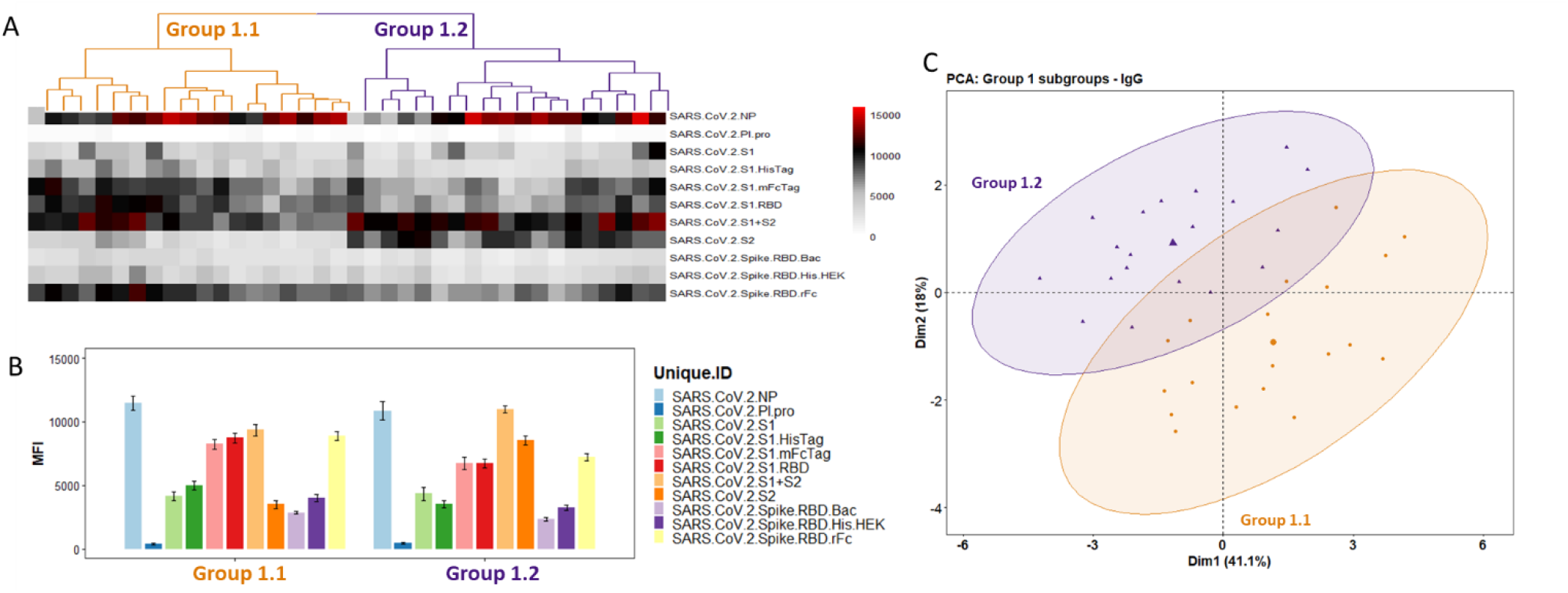
Group 1 Cluster Analysis and PCA demonstrates two subgroups. Panel A, heatmap showing the reactivity to the SARS-CoV-2 antigens. Samples were clustered using the hierarchical clustering analysis. The dendrogram was cut (and color coded) to a final cluster number equal 3. On B, a bar graph showing the mean reactivity and the standard error of each cluster to each individual SARS-CoV-2 antigen. Panel C, PCA analysis showing the spatial distribution of the samples classified as 1.1 and 1.2 for the first and second principal components.

**Figure 5.**
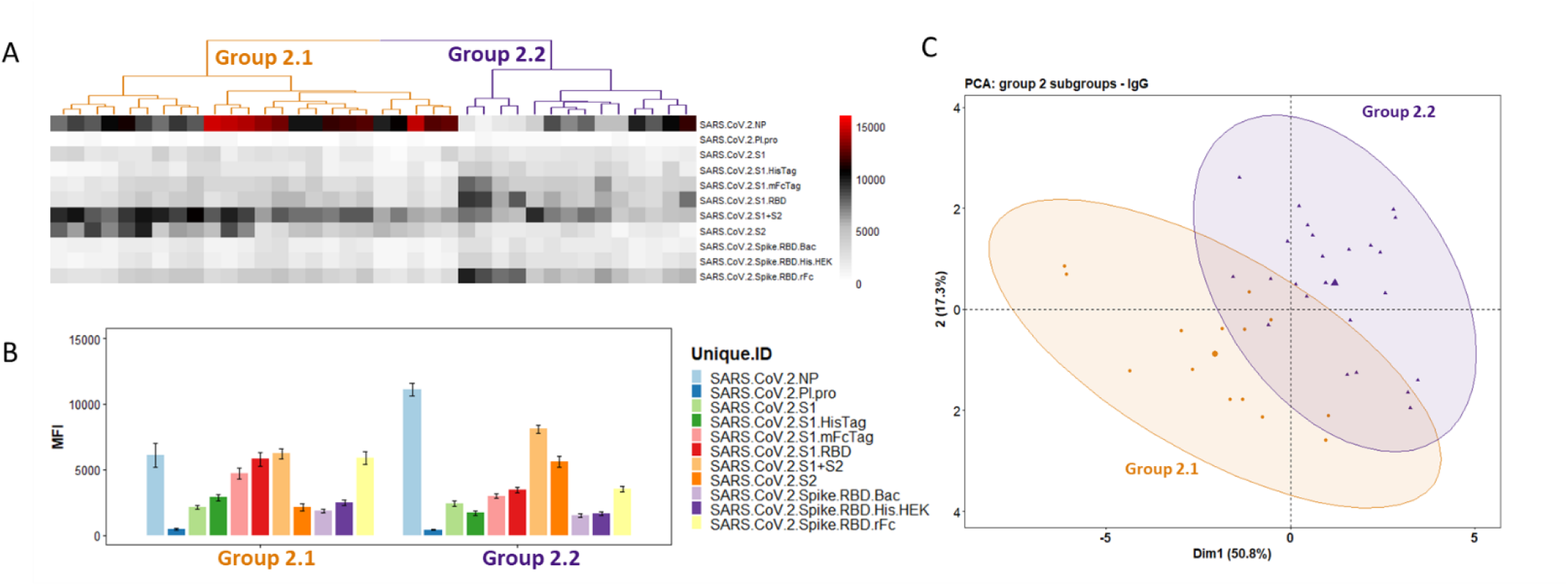
Group 2 Cluster Analysis and PCA demonstrates two subgroups. Panel A, heatmap showing the reactivity to the SARS-CoV-2 antigens. Samples were clustered using the hierarchical clustering analysis. The dendrogram was cut (and color coded) to a final cluster number equal 3. On B, a bar graph showing the mean reactivity and the standard error of each cluster to each individual SARS-CoV-2 antigen. Panel C, PCA analysis showing the spatial distribution of the samples categorized as subgroups 2.1 and 2.2 for the first and second principal components.

**Figure 6.**
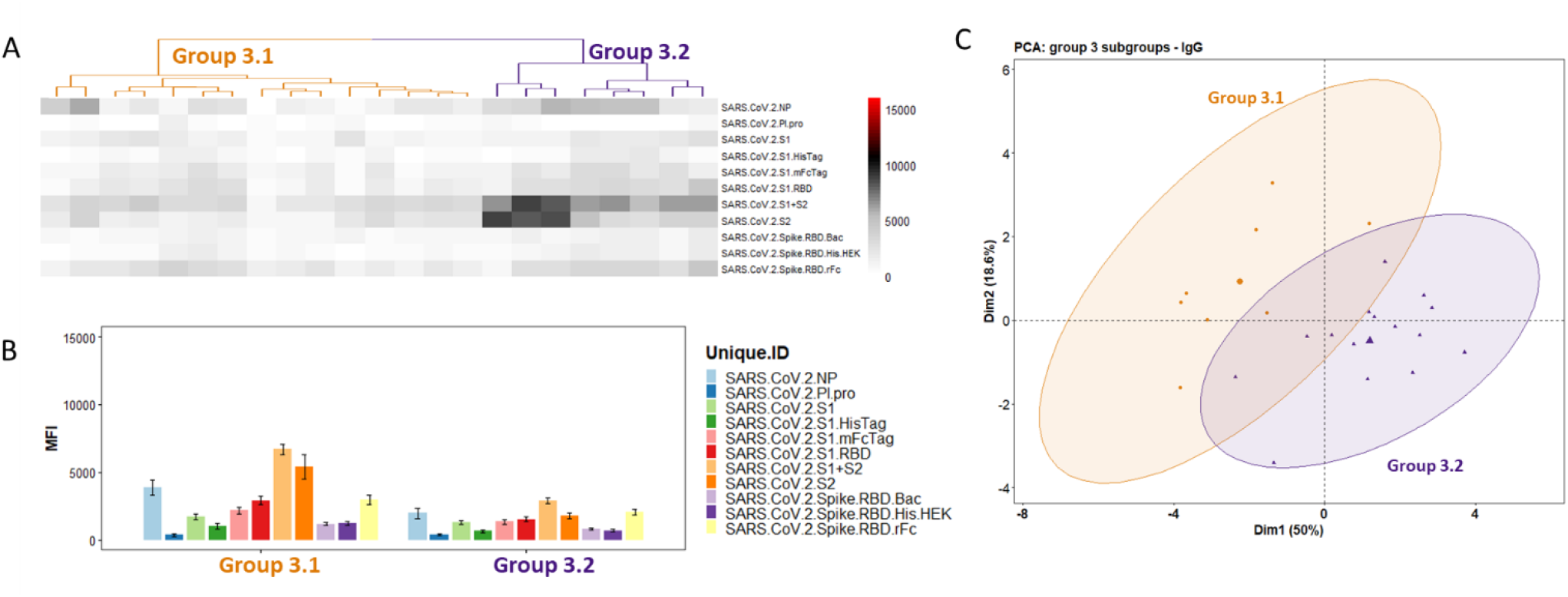
Group 3 Cluster Analysis and PCA demonstrates two subgroups. Panel A, heatmap showing the reactivity to the SARS-CoV-2 antigens. Samples were clustered using the hierarchical clustering analysis. The dendrogram was cut (and color coded) to a final cluster number equal 3. On B, a bar graph showing the mean reactivity and the standard error of each cluster to each individual SARS-CoV-2 antigen. Panel C, PCA analysis showing the spatial distribution of the samples categorized as subgroups 3.1 and 3.2 for the first and second principal components.

### Multiplex antigen classification of CCP

The efficacy of CCP may vary depending on the donor, and there are different ways to classify the plasma before it is administered. One criterion is whether the donor was a PCR+ confirmed case. Factors such as clinical disease severity and the time from symptom onset or recovery to collection of CCP units may contribute to the classification. Antibody level is a measurement that can be used to qualify CCP and antibodies against different antigen targets including the total spike S1/S2, S1, S2, RBD and NP proteins can be considered. Virus neutralization assay titers are another metric and pseudoviruses have been developed for this purpose to make it more convenient than live SARS-CoV-2 virus neutralization assays for routine use. In order to use these quantitative measurements to qualify CCP for clinical administration, a threshold can be established, to qualify a unit of plasma as acceptable for clinical use or rejected.

A multiplex antigen classification of CCP can also be considered as a criterion to qualify and accept or reject donor plasma for transfusion to COVID-19 patients. For example, all of the highly reactive COVAM Group 1 specimens **(Figure 3**) could be accepted and all of the low reactive Group 3 specimens rejected. Or subsets of any of the groups (**Figure 4-6**) could be accepted or rejected based on knowledge of the clinical efficacy of each major group or subgroup The results plotted in **Figure 7** show how multiplex antigen classification differs from criteria based on changing the binary threshold of a single assay. In **Figure 7A** the RVPN titer threshold was moved from the canonical standard 40, to 160, 320 and 640. Group 3 specimens increasingly fall below the cutoff as the threshold increases. Group 2 specimens are mostly above threshold at the 160 threshold and increasingly fall below the threshold as it is increased. Interestingly, all but one of the Group 1 specimens are below the highest RVPNT threshold of 640.

**Figure 7.**
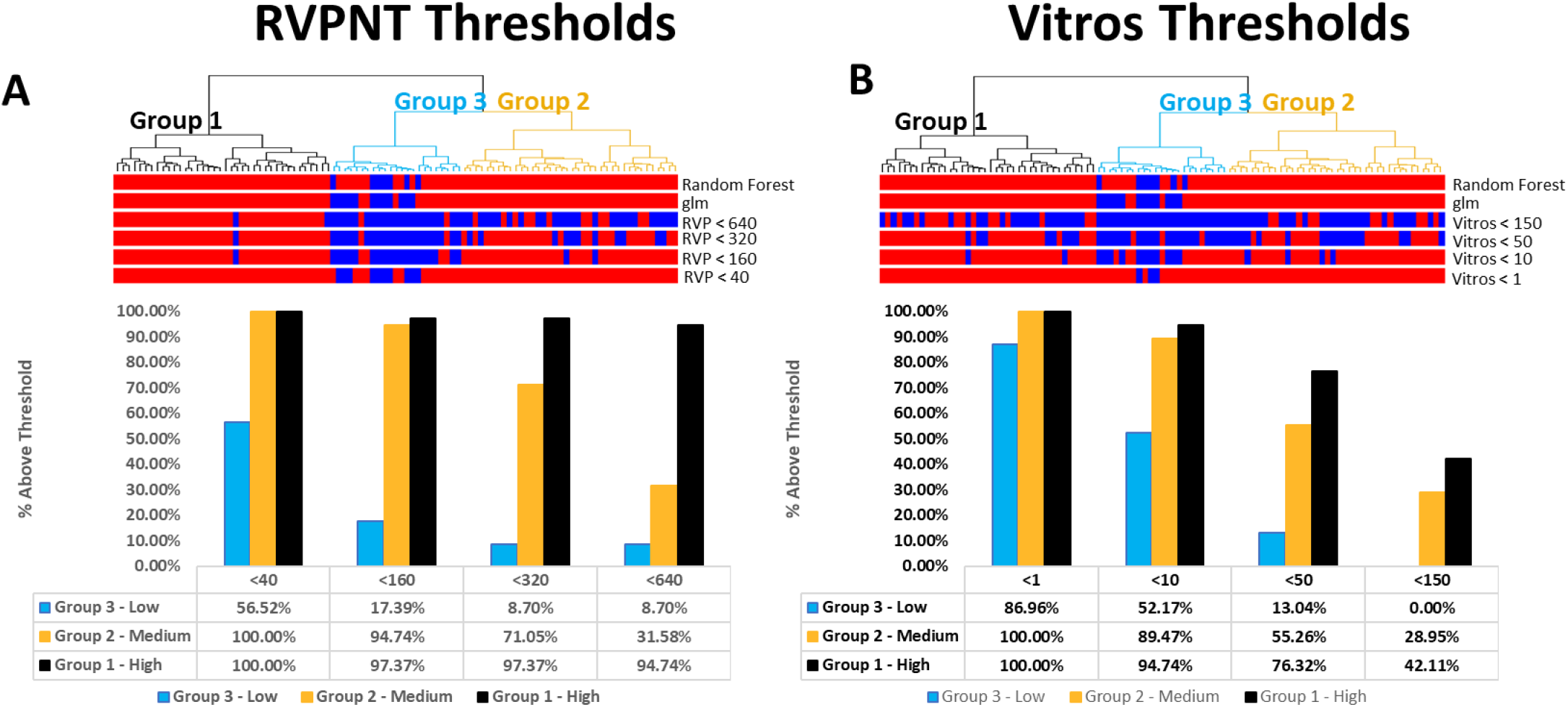
SARS-CoV-2 Reactivity Classification. In both Panels A (comparison to RVPNT NT50 thresholds) and B (comparison to VITROS CoV2T S/C thresholds), the bars across the top represent the classification for each given sample. The color red represents samples classified as reactive and the color blue, samples classified as non-reactive. The top 2 bars represent the COVAM reactivity predictions with the top bars the prediction based on the Random Forest model and the second bars the prediction from the logistic regression model. The bar graphs represent the % of samples classified as reactive on each cutoff value.

The VITROS CoV2T assay threshold can also be adjusted to select the specimens with higher signals (**Figure 7B**). Moving the S/C threshold from 1 to 10 preferentially removes a high proportion of the low-level reactive Group 3 specimens. At the highest threshold of 150 all of the Group 3 specimens are removed while 40% and 30% of the Group 1 and 2 specimens, respectively, are above the threshold.

These assay-dependent and threshold-dependent differences in the classification of plasma specimens is further evident from the results in **Figure 8**. Here the group 1, 2 and 3 specimens were separately sorted by the RVPNT titer (**Figure 8A**). The Titer Cutoff lines on **Figure 8A** shows how most of the Group 1 samples are above the threshold even at the highest 640 titer. The Group 2 and Group 3 specimen classifications are affected more by increasing the Cutoff titer. Similarly, the VITROS CoV2T data is plotted in **Figure 8B** and 3 different assay cutoff values are shown. It is evident that the VITROS CoV2T intensity is not well correlated to the RVPNT titer, and scatterplots (not show) give R^2^ = 0.037. Increasing the VITROS CoV2T threshold cuts through all the groups.

**Figure 8.**
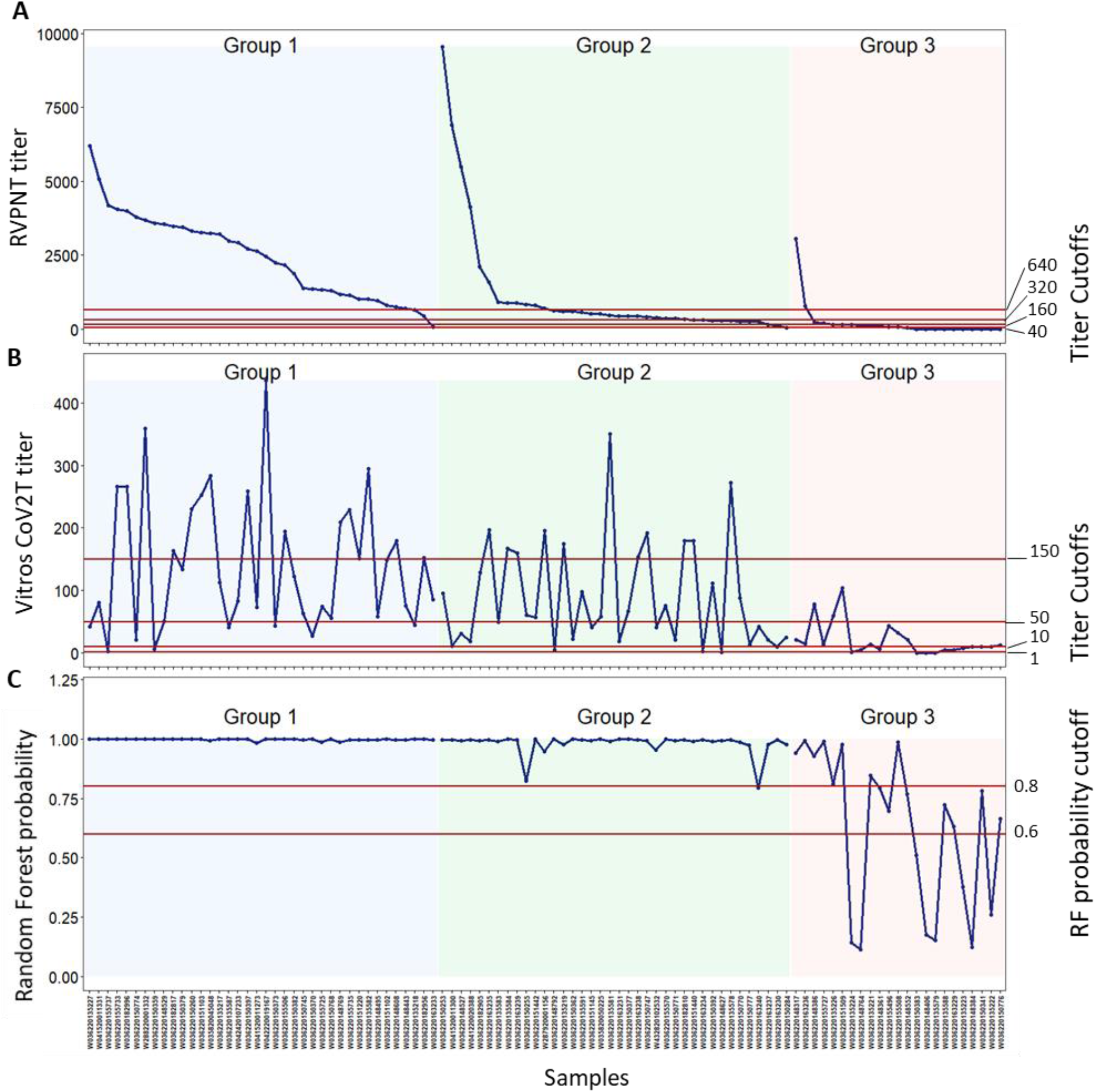
SARS-CoV-2 RVPNT and VITROS-CoV2T titers. In both Panels A (RVPNT NT50 titers) and B (VITROS CoV2T S/C titers), the samples were sorted by their cluster classification and the RVPNT titers. The red, horizontal lines represent different cutoffs being, for RVPNT 1:40; 1:160; 1:320 and 1:640; for VITROS CoV2T S/C > 1, 10, 50 and 150 and for the Random Forest prediction probability 0.6 and 0.8.

These results indicate that RVPNT titer, VITROS CoV2T titer, and the COVAM multiplex cluster analysis patterns select different collections of plasma specimens.

## Discussion

Passive antibody transfer using convalescent plasma has previously been used to treat infectious diseases that involve the respiratory system including influenza [4-7]. Prior experience in epidemics with convalescent plasma (CP) containing antibodies to viruses has demonstrated variable indications of therapeutic efficacy against Influenza, Argentine Hemorrhagic Fever, and SARS. Characterizing antibody titers in plasma to viruses has indicated a correlation with therapeutic efficacy. An Expanded Access Protocol (EAP) clinical trial led by the Mayo Clinic has been completed resulting in >65,000 units of apheresis plasma collected from convalescent COVID-19 patients and treating >35,000 acutely infected COVID-19 in-patients. Anti-SARS-CoV-2 antibody levels in donor plasma are variable, and a small statistically significant (p<0.05) improvement in the clinical outcome was reported in a subset of patients who were treated early after symptom onset and received the plasmas with higher Ab titers compared to patients who received plasma later in the disease course. The results suggest that the SARS-CoV-2 specific Ab content of CCP may be associated with better clinical outcomes in the appropriate patient subset. Definitive safety and efficacy of CCP for the treatment of COVID-19 is awaiting completion of placebo controlled, prospective and randomized clinical trials (RCTs).

Clinical serodiagnostic tests usually measure antibody levels against a single antigen or epitope and a single Ig isotype. The output of current EUA SARS-CoV-2 Ab assays are a binary positive/negative, above or below a threshold with or without reporting intensity of reactivity levels above the threshold. The assay cutoff threshold is determined by probing a well-characterized collection of positive and negative control samples. Some tests of this type mix multiple antigens together and use secondary Abs that recognize multiple isotypes to produce an aggregate value with a binary positive/negative result.

Methodologies like COVAM that quantitatively interrogate multiple specific signal intensity levels independently to classify specimens were called “chemometrics” in the pre-genomic period. Modern multivariant statistical approaches that interrogate and classify a “fingerprint” of individual genomic measurements have been developed to classify genome sequence and expression data, and these computational methods can also be applied to clinical diagnostics. The test results from the COVAM assay described here take advantage of multivariant machine learning, pattern recognition, and clustering to classify plasma samples that have many antibodies at different levels against numerous SARS-CoV-2 antigens.

The COVAM was constructed to independently measure immunoglobulin isotype and subtype levels in serum or plasma samples against each of the individual antigens printed on the array. We use two computational approaches, either a generalized linear model (glm) or a Random Forest (RF) prediction model to classify individual specimens as either Reactive or Non-Reactive against the SARS-CoV-2 antigens. We use a training set of 88 PreCoV specimens collected in July-September 2019 and positive specimens from PCR+ confirmed COVID-19 cases. We evaluated 99 coded CCP plasma samples and produced binary positive/negative results that were 96% concordant with the FDA EUA Ortho VITROS CoV2T immunoassay and 93% with the RVPNT assay. All of the discordances were weak responders around the threshold of the comparators.

Although the binary classification of COVAM, VITROS CoV2T and RVPNT assays are highly concordant, multivariant COVAM analysis of quantitative results reveals distinct differences among the 99 CCP specimens analyzed in this study. Unsupervised K-means analysis, as well as hierarchical clustering reveal 3 main clusters with distinct reactivity intensities and patterns. The dendrogram analysis reveals at least 2 additional subgroups within each of the major groups that separate based on reactivity patterns against the antigens on the array. We could not recapitulate these group classifications by simply moving the reactivity thresholds of either the VITROS CoV2T or RVPNT assays.

The variable reactivity patterns between individuals observed in this study is suggestive of polyclonal immune responses with different antibody specificities which should be examined in light of clinical outcomes in recipients of CCP transfusions; if predictive the COVAM assay could be employed to qualify CCP for administration as a treatment for acute COVID. The use of a SARS-CoV-2 antigen array of this kind could also be useful to interrogate vaccine performance in preclinical and clinical studies in order to understand Ab responses associated with protection from acquisition of infection, and prediction of clinical severity of disease in vaccine breakthrough infections. An analysis to identify antibody patterns associated with treatment efficacy can be done retrospectively on aliquots from plasma that have already been used for treatment in ongoing clinical trials of CCP and vaccines.

## Acknowledgements

This work was supported by the HDTRA1-16-C-0009, HDTRA1-18-1-0035, HDTRA1-18-1-0036 and a University of California, Irvine CRAFT-COVID grants.The findings and conclusions in this report are those of the authors and do not necessarily represent the official position or policy of the funding agencies and no official endorsements should be inferred.

## Author Approvals

All authors have seen and approved this manuscript. This manuscript is an original work and has not been accepted to or published elsewhere.

