## Supplementary Table 1 for "Distinct SARS-CoV-2 Antibody Reactivity Patterns in Coronavirus Convalescent Plasma Revealed by a Coronavirus Antigen Microarray"

COVAM antigens in order displayed on the figures. All antigens were supplied by Sino Biological Inc.

| Antigen ID | Virus | Strain | Protein | Catalogue No |
|---|---|---|---|---|
| SARS.CoV.2.NP | SARS-CoV-2 | SARS-CoV-2 | NP | 40588-V08B |
| SARS.CoV.2.Pl.pro | SARS-CoV-2 | SARS-CoV-2 | Papain-like protease | 40593-V07E |
| SARS.CoV.2.S1 | SARS-CoV-2 | SARS-CoV-2 | S1 subunit | 40591-V08B1 |
| SARS.CoV.2.S1.HisTag | SARS-CoV-2 | SARS-CoV-2 | S1, (His Tag) | 40591-V08H |
| SARS.CoV.2.S1.mFcTag | SARS-CoV-2 | SARS-CoV-2 | S1, (mFc Tag) | 40591-V05H1 |
| SARS.CoV.2.S1.RBD | SARS-CoV-2 | SARS-CoV-2 | S1-RBD | 40592-V05H |
| SARS.CoV.2.S1+S2 | SARS-CoV-2 | SARS-CoV-2 | S1+S2 | 40589-V08B1 |
| SARS.CoV.2.S2 | SARS-CoV-2 | SARS-CoV-2 | S2 | 40590-V08B |
| SARS.CoV.2.Spike.RBD.Bac | SARS-CoV-2 | SARS-CoV-2 | RBD | 40592-V08B |
| SARS.CoV.2.Spike.RBD.His.HEK | SARS-CoV-2 | SARS-CoV-2 | RBD | 40592-V08H |
| SARS.CoV.2.Spike.RBD.rFc | SARS-CoV-2 | SARS-CoV-2 | RBD | 40592-V31H |
| SARS.CoV_NP | SARS-CoV-2 | SARS | NP | 40143-V08B |
| SARS.CoV_PLpro | SARS-CoV-2 | SARS | PLpro | 40524-V08E |
| SARS.CoV_S1.HisTag | SARS-CoV-2 | SARS | S1, (His Tag) | 40150-V08B1 |
| SARS.CoV_S1.RBD.HisTag | SARS-CoV-2 | SARS | S1-RBD, (His Tag) | 40150-V08B2 |
| SARS.CoV_S1.RBD.rFcTag | SARS-CoV-2 | SARS | S1-RBD, rFc Tag | 40150-V31B2 |
| MERS.CoV_NP | MERS | MERS | NP | 40068-V08B |
| MERS.CoV_S1.AA1.725.His.HEK | MERS | MERS | S1, N-(AA1-725, His Tag)_A | 40069-V08H |
| MERS.CoV_S1.RBD.367.606.rFcTag | MERS | MERS | S1-RBD, N-(AA367-606, rFc Tag) | 40071-V31B1 |
| MERS.CoV_S1.RBD.383.502.mFcTag | MERS | MERS | S1-RBD, N-(AA383-502, mFc Tag) | 40071-V05B |
| MERS.CoV_S2 | MERS | MERS | S2 | 40070-V08B |
| DcCoV.HKU23.NP | CommonCoV | HKU23-368F | NP | 40458-V08B |
| hCoV.229E.S1 | CommonCoV | 229E | S1 | UN1-2 |

| | | | | |
|---|---|---|---|---|
| hCoV.229E.S1_S2 | CommonCoV | 229E | S1+S2 | UN2-2 |
| hCoV.HKU1.HE | CommonCoV | HKU1 | HE | UN1-4-1 |
| hCoV.HKU1.S1_AA1.760 | CommonCoV | HKU1 | S1, N-(AA1-760) | 40021-V08H |
| hCoV.HKU1.S1_AA13.756 | CommonCoV | HKU1 | S1, N-(AA13-756) | UN1-3 |
| hCoV.HKU1.S1_S2 | CommonCoV | HKU1 | S1+S2 | UN2-3 |
| hCoV.NL63.S1 | CommonCoV | NL63 | S1 | UN1-1 |
| hCoV.NL63.S1_S2 | CommonCoV | NL63 | S1+S2 | UN2-1 |
| hCoV.OC43.HE | CommonCoV | OC43 | HE | UN1-6 |
| hCoV.OC43.S1 | CommonCoV | OC43 | S1 | UN1-5 |
| hCoV.OC43.S1_S2 | CommonCoV | OC43 | S1+S2 | UN2-4 |
| Flu.B_Mal/.HA1 | Influenza | B/Malaysia/2506/2004 | HA1 | 11716-V08H1 |
| Flu.B_Mal/.HA1+HA2 | Influenza | B/Malaysia/2506/2004 | HA1+HA2 | 11716-V08H |
| Flu.B_Phu/.HA1 | Influenza | B/Phuket/3073/2013 | HA1 | 40498-V08H1 |
| Flu.B_Phu/.HA1+HA2 | Influenza | B/Phuket/3073/2013 | HA1+HA2 | 40498-V08B |
| Flu.H1N1.HA1 | Influenza | A/Beijing/22808/2009 | HA1 | 40035-V08H1 |
| Flu.H1N1.HA1+HA2 | Influenza | A/Beijing/22808/2009 | HA1+HA2 | 40035-V08H |
| Flu.H3N2.HA1 | Influenza | A/Texas/50/2012 | HA1 | 40354-V08H1 |
| Flu.H3N2.HA1+HA2 | Influenza | A/Texas/50/2012 | HA1+HA2 | 40354-V08B |
| Flu.H5N1.HA1 | Influenza | A/Vietnam/1203/2004 | HA1 | 10003-V06H1 |
| Flu.H5N1.HA1+HA2 | Influenza | A/Vietnam/1203/2004 | HA1+HA2 | 10003-V06H3 |
| Flu.H7N9.HA1 | Influenza | A/Anhui/1/2013 | HA1 | 40103-V08H1 |
| Flu.H7N9.HA1+HA2 | Influenza | A/Anhui/1/2013 | HA1+HA2 | 40103-V08H |
| hAdV3.Fiber | Adenovirus | hAdV-3/45659 | Fiber | UN5-9 |
| hAdV3.Penton | Adenovirus | hAdV-3/45659 | Penton | UN5-10 |
| hAdV4.Fiber | Adenovirus | hAdV-4/28280 | Fiber | UN5-14 |

| | | | | |
|---|---|---|---|---|
| hAdV4.Penton | Adenovirus | hAdV-4/28280 | Penton | UN5-15 |
| hMPV.A_G.52N.228N | hMPV | PER/CFI0320/2010/A Glycoprotein | hMPV-A_G (52N-228N) | UN6-3 |
| hMPV.B_F.280D.490G | hMPV | PER/CFI0320/2010/A Glycoprotein | hMPV-B_F (280D-490G) | UN6-5 |
| hMPV.B_G.52D.238S | hMPV | PER/CFI0466/2010/B Glycoprotein | hMPV-B_G (52D-238S) | UN6-6 |
| hPIV.1.12O3_F | Parainfluenza | hPIV-1 12O3 | F | UN5-1 |
| hPIV.1.12O3_H | Parainfluenza | hPIV-1 12O3 | H | UN5-2 |
| hPIV.3.2010_H | Parainfluenza | hPIV-3 USA/10991B/2010 | H | UN5-6 |
| hPIV.4.b.2016_H | Parainfluenza | hPIV-4b/10-H2/2016 | H | UN5-8 |
| RSV.A.F | RSV | LA2-94/2013 | F | UN3-1 |
| RSV.A.G | RSV | LA2-94/2013 | G | UN4-1 |
| RSV.B.F | RSV | TH-10526/2014 | F | UN3-2 |
| RSV.B.G | RSV | B1 | G | 13029-V08H |
